# Exploration of American Elderberry Bioactive Compounds for Inhibition of the Influenza Virus Polymerase Acidic Endonuclease

**DOI:** 10.1101/2025.11.12.688002

**Authors:** Amanda Dwikarina, Hsin-Yeh Hsieh, Michael C. Greenlief, Andrew L. Thomas, Chung-Ho Lin

## Abstract

Elderberries have been commonly used to relieve symptoms of influenza and the common cold. In this study, the antiviral activity of juices of American elderberry (*Sambucus nigra* subsp. *canadensis*) cultivars were evaluated against the recombinant N-terminal domain of *Influenza* A Polymerase Acidic protein using a Fluorescence Resonance Energy Transfer (FRET)-based endonuclease assay. The American elderberry cultivar Ozark showed the strongest inhibitory activity among 21 cultivars, followed by the wild accession 1196. The juice from four American elderberry cultivars Ozark, accession 1892, 1196, and 2084 demonstrated stronger inhibitory effect against nPA protein (>65% inhibition) compared to the European elderberry cultivar Haschberg, which showed 62% inhibition, suggesting a higher abundance of bioactive compounds in these cultivars, capable of interfering with *Influenza* PA activity. Among the elderberry compounds tested, gallic acid (IC_50_ 11.56 ± 1.80 µM), myricetin (IC_50_ 16.02 ± 0.06 µM), caffeic acid (IC_50_ 22.75 ± 0.9 µM), and luteolin (IC_50_ 27.92 ± 0.72 µM), showed the strongest inhibition of nPA endonuclease activity. These findings support the application of American elderberry in nutraceutical product formulations designed to help manage flu symptoms. To the best of our knowledge, this is the first study elucidating the anti-influenza mode of action of American elderberry against influenza molecular targets.

## 1. Introduction

*Influenza* is a respiratory pathogen that causes the seasonal flu. Influenza could lead to significant morbidity and mortality worldwide. Although vaccines and antivirals are available, both are hindered by limitations, including restricted treatment windows and the emergence of resistance mutations. These challenges underscore the urgent need for alternative or complementary antiviral strategies that utilize viral targets.

There is a strong societal trend towards the use of natural health products to manage respiratory infections. Elderberry-based supplements, particularly syrups and juices, are widely marketed to alleviate cold and Influenza-like symptoms, reflecting consumer preferences for natural remedies. Previous studies on European elderberry (*Sambucus nigra*) demonstrated the benefit of elderberry consumption on health. A clinical study showed that consumption of elderberry syrup containing elderberry extract reduced the duration and severity of Influenza-like symptoms in adults. ^1,2^. Previously, the studies with *in vitro* cell culture reported that elderberry extract not only suppresses *Influenza* replication but also modulates host immune responses, such as enhanced inflammatory cytokine production in host cells ^3–5^.

The anti-*Influenza* activity of elderberries has become the subject of study, with a primary focus on European elderberries ^6–9^. Several antiviral mechanisms of elderberry have been proposed. A previous study utilizing plaque reduction assays demonstrated that the juice of European elderberry (*Sambucus nigra*) showed antiviral effects against *Influenza* by directly interacting with viral surface glycoproteins, hemagglutinin and neuraminidase, hence blocking viral entry ^5^. Pretreatment of viruses with elderberry juice before being exposed to the cells reduced the *Influenza* infection, while extended exposure of elderberry juice to the cells after *Influenza* infection significantly enhanced antiviral activity, suggesting that elderberry interferes with multiple stages of the viral life cycle. However, the molecular targets underlying antiviral activity on viral replication remain poorly characterized. Moreover, the antiviral properties of American elderberry (*Sambucus nigra* subsp. *canadensis*) have not been well described in comparison to its European relatives.

Based on previous studies, elderberry has been suggested to alleviate the symptoms of influenza, raising the possibility that its bioactive compounds may act by targeting early stages of infection and interfering with the viral transcription mechanism within host cells. During *Influenza* infection, viral mRNA requires both a 5’cap structure and a poly(A) tail to be recognized and transcribed by the host transcriptional machinery. The process to acquire them is mediated by the *Influenza* RNA-dependent RNA polymerase (RdRp) complex. The viral RdRp complex consists of three subunits, Polymerase Basic 1 (PB1), Polymerase Basic 2 (PB2), and Polymerase Acidic (PA) protein. The PA protein is an essential component of the viral RdRp complex, due to the endonuclease active site in its N-terminal domain that is necessary for “cap-snatching” from the hosts’ pre-mRNAs to initiate viral transcription ^10^. In this mechanism, PB2 would bind to the 5’ cap of the host pre-mRNA molecules for PA endonuclease to cleave it off from the host pre-mRNA, and then PB1 has polymerase activity to add it on the viral RNA and generate a capped primer required for initiating transcription in the host cells. The PB1 is also responsible for the addition of a poly(A)tail. The clinical success of baloxavir marboxil, a selective inhibitor of the PA endonuclease, validated this complex as a target for antivirals. Baloxavir binds to the active site of the PA protein, preventing it from cleaving the host RNA ^11^. By blocking the cap-snatching process, baloxavir inhibits virus’s ability to replicate and produce new viral particles.

Flavonoids, a diverse group of plant-derived specialized metabolites that have well-documented antiviral activity, are abundant in elderberries. It represents a promising source of diverse small molecules with the potential to inhibit PA endonuclease activity. This study evaluated the antiviral potential of the elderberry juices and bioactive compounds identified in American elderberry against the PA protein of *Influenza* A using a FRET-based endonuclease assay to explore critical bioactive compounds underlying the antiviral effects of elderberry.

## 2. Materials and Methods

### 2.1 Materials

The cDNA template encoding Influenza A/California/07/2009 (H1N1) PA ORF was purchased from Integrated DNA Technologies (Coralville, IA, USA). StrataClone PCR cloning kit was purchased from Agilent Technologies (La Jolla, CA, USA). The *Escherichia coli* expression host BL21(DE3)RIL competent cells were obtained from Agilent Technologies (La Jolla, CA, USA). Terrific broth expression media, Ni-NTA His-spin columns, and HEPES buffer were purchased from Thermo Fisher Scientific (Waltham, MA, USA). Purification buffers (binding buffer, wash buffer, and elution buffer) were purchased from Zymo (Irvine, CA, USA). Chemical standards of bioactive compounds in American elderberry were purchased from Sigma-Aldrich (St. Louis, MO, USA) but isorhamnetin 3-rutinoside was purchased from Fisher Chemical (Fair lawn, NJ, USA) with purity >95% (Table 1).

**Table 1.** The half maximal inhibitory concentration (IC_50_) values of bioactive compounds against *Influenza* nPA protein.

| No. | Compound Name | Sources | Purity |
| --- | --- | --- | --- |
| 1 | Caffeic acid | Sigma-Aldrich | >98% |
| 2 | Cryptochlorogenic acid | Sigma-Aldrich | >98% |
| 3 | Catechin hydrate | Sigma-Aldrich | >98% |
| 4 | Chlorogenic acid | Sigma-Aldrich | >95% |
| 5 | p-Coumaric acid | Sigma-Aldrich | >98% |
| 6 | Cyanidin 3,5-O-diglucoside | Sigma-Aldrich | >98% |
| 7 | Cyanidin 3-O-galactoside | Sigma-Aldrich | >95% |
| 8 | Cyanidin 3-O-glucoside | Sigma-Aldrich | >98% |
| 9 | Cyanidin 3-O-rutinoside | Sigma-Aldrich | >90% |
| 10 | Cyanidin 3-O-sambubioside | Sigma-Aldrich | >95% |
| 11 | Cyanidin 3-O-sophoroside | Sigma-Aldrich | >95% |
| 12 | Delphinidin 3-O-rutinoside | Sigma-Aldrich | >95% |
| 13 | 3-4, Dihydroxybenzoic acid | Sigma-Aldrich | >95% |
| 14 | Epicatechin | Sigma-Aldrich | >98% |
| 15 | Ferulic acid | Sigma-Aldrich | certified reference standard |
| 16 | Gallic acid | Sigma-Aldrich | >98% |
| 17 | Hyperoside (quercetin 3-galactoside) | Sigma-Aldrich | >95% |
| 18 | Isorhamnetin | Sigma-Aldrich | >95% |
| 19 | Isorhamnetin 3-rutinoside | Fisher Chemical | >95% |
| 20 | Kaempferol | Sigma-Aldrich | >97% |
| 21 | Kaempferol-3-glucoside | Sigma-Aldrich | >95% |
| 22 | Kaempferol 3-O-rutinoside | Sigma-Aldrich | >95% |
| 23 | Luteolin | Sigma-Aldrich | >95% |
| 24 | Myricetin | Sigma-Aldrich | >96% |
| 25 | Naringenin | Sigma-Aldrich | >95% |
| 26 | Neochlorogenic acid | Sigma-Aldrich | >98% |
| 27 | Pelargonidin 3-O-glucoside | Sigma-Aldrich | >95% |
| 28 | Peonidin 3-O-glucoside | Sigma-Aldrich | >90% |
| 29 | Quercetin | Sigma-Aldrich | >95% |
| 30 | Isoquercitrin (quercetin-3-glucoside) | Sigma-Aldrich | >90% |
| 31 | Rutin (Quercetin-3-rutinoside) | Sigma-Aldrich | >95% |
| 32 | Isorhamnetin 3-O-glucoside | Sigma-Aldrich | >98% |

**Table 1.** List of 32 bioactive compounds identified in American elderberry juices that are used in this study.

| No. | Compound Name | IC<sub>50</sub> (μM) |  |
| --- | --- | --- | --- |
|  |  | AVG | SEM |
| 1 | Caffeic acid | 22.75 | 0.90 |
| 2 | Cryptochlorogenic acid | NI | NI |
| 3 | Catechin hydrate | NI | NI |
| 4 | Chlorogenic acid | 33.28 | 2.63 |
| 5 | p-Coumaric acid | NI | NI |
| 6 | Cyanidin 3,5-O-diglucoside | NI | NI |
| 7 | Cyanidin 3-O-galactoside | NI | NI |
| 8 | Cyanidin 3-O-glucoside | NI | NI |
| 9 | Cyanidin 3-O-rutinoside | 93.64 | 23.94 |
| 10 | Cyanidin 3-O-sambubioside | 120.60 | 28.02 |
| 11 | Cyanidin 3-O-sophoroside | NI | NI |
| 12 | Delphinidin 3-O-rutinoside | 71.11 | 12.83 |
| 13 | 3-4, Dihydroxybenzoic acid | NI | NI |
| 14 | Epicatechin | NI | NI |
| 15 | Ferulic acid | NI | NI |
| 16 | Gallic acid | 11.56 | 1.81 |
| 17 | Hyperoside (quercetin 3-galactoside) | NI | NI |
| 18 | Isorhamnetin | NI | NI |
| 19 | Isorhamnetin 3-rutinoside | NI | NI |
| 20 | Kaempferol | NI | NI |
| 21 | Kaempferol-3-glucoside | NI | NI |
| 22 | Kaempferol 3-O-rutinoside | NI | NI |
| 23 | Luteolin | 27.92 | 0.72 |
| 24 | Myricetin | 16.01 | 0.06 |
| 25 | Naringenin | NI | NI |
| 26 | Neochlorogenic acid | 49.67 | 4.45 |
| 27 | Pelargonidin 3-O-glucoside | NI | NI |
| 28 | Peonidin 3-O-glucoside | 104.75 | 1.04 |
| 29 | Quercetin | 39.80 | 3.79 |
| 30 | Quercetin-3-glucoside (isoquercitrin) | NI | NI |
| 31 | Rutin (Quercetin-3-rutinoside) | NI | NI |
| 32 | Isorhamnetin 3-O-glucoside | NI | NI |

\* NI = No inhibition
\*\* Data presented as mean value ± SEM (n=2)

### 2.2 Plant materials

The American elderberry juices subjected to this study were previously described in Dwikarina *et al*. (2024). In brief, American elderberry (*Sambucus nigra* subsp. *canadensis*) fruits from 18 propagated accessions and three established cultivars (Tabel S1) were harvested from plantings at the University of Missouri’s Southwest Research, Extension, and Education Center near Mt. Vernon, Missouri, USA. The fruits were harvested at peak ripeness and immediately frozen. The fruit was then de-stemmed, thawed at room temperature, and the juice was pressed by hand. It was filtered through a kitchen sieve, aliquoted into 50 mL polypropylene tubes, and re-frozen at -20 °C until further analysis.

The fruits of the European elderberry (*Sambucus nigra*) were of cultivar Haschberg. Theseedlings were originally purchased from a local nursery in Portland, Oregon (https://onegreenworld.com/) and cultivated under greenhouse conditions at the University of Missouri, Columbia, MO, USA. The fruits were harvested at full ripeness during the summer of 2024 and immediately frozen. Prior to processing, fruits were de-stemmed, thawed at room temperature, and placed in 15 ml tubes for juice extraction. Berries were gently pressed with a clean spatula to extract juice, which was then centrifuged at 4700 rpm for 10 minutes at 10 °C to separate the pomace. The supernatant was then transferred to a 2 mL Eppendorf tube and stored at -20°C until further analysis.

### 2.3 Gene cloning

PCR amplification of the gene encoding the N-terminal domain of Influenza A/California/07/2009 (H1N1) PA ORF, corresponding to amino acids residues 1-257, was performed with annealing temperature of 53.6 ^°^C and 55.5 ^°^C, and extension time for 1 min at 72 ^°^C. Forward primer sequences containing *XbaI* restriction site (in bold) that were used for PCR in this study were: 5’ **TCT AGA** ATG GAA GAC TTT GTG CCGA CAA TG 3’. Reverse primer sequences containing *XhoI* restriction site (in bold) that were used for PCR in this study were: 5’ **CTC GAG** AAT TTT GGC GTT CAC TTC TTT TG 3’.

The *npa* gene fragments were then cloned into PCR cloning vector *p*SCA (Agilent Technologies, La Jolla, CA, USA). *E. coli* SoloPack cells were then transformed with *npa* cloned *p*SCA using heat shock method by incubation at 37^°^C for 5 min and then revivable incubation at 37^°^C for 1 hour after adding 250 *μ*l of LBroth, for cloning amplification. Gene fragments of *npa* were dropped out of the cloned *p*SCA by the dual digestion of *XbaI* and *XhoI* restriction enzymes (New England Biolabs, Ipswich, MA) and subcloned to a T7 expression vector, *p*ET303 vector plasmid (Invitrogen) in frame with a nucleotide region encoding for a C-terminal 6X-HisTag. The ligation reaction was prepared by mixing the digested vector and the *npa* fragements with T4 DNA ligase (NEB England Biolabs, Ipswich, MA) following the manufacturer’s protocol. The ligation reactions were incubated at 16^°^C overnight. The ligation products were then transferred into *E. coli* competent cells strain DH5α and were inoculated into LB agar media containing 100 µg/mL ampicillin. DNA extraction was performed to purified recombinant plasmids using Wizard *PLUS SV* minipreps DNA purification kit (Promega, Madison, WI, USA). The purified recombinant plasmids were digested by *XbaI* and *XhoI* restriction enzymes and run on the 1% agarose gel containing ethidium bromide. The insertion of the *nPA* gene fragments into *pET303* was confirmed by size, then the clones were sent for sequence verification at MU DNA core facility. The *p*ET303 carrying the nPA ORF was transferred into the T7 expression host *BL21(DE3)RIL* strain for protein expression.

### 2.4 Protein expression and purification

The nPA protein was expressed and purified following as described previously ^12^ with modification. Overnight cultures were grown in 5 mL of LB broth containing 100 µg/mL ampicillin and 10 µg/mL chloramphenicol and incubated with 200 rpm agitation at 37°C overnight. This starter culture was then used to inoculate 1 L of expression media (Terrific Broth media with 0.2% dextrose, 0.1 mM MnCl_2,_ and 0.1 mM MgSO_4_) containing 100 µg/ml ampicillin and 10 µg/ml chloramphenicol. The culture was incubated at 37°C with shaking at 180 rpm until an optical density at 600 nm of 0.8 was reached. Expression of nPA protein was then induced by adding IPTG to a final concentration of 1 mM. The induced culture was incubated at 22^°^C with agitation at 180 rpm for 16-18 hours. The culture for protein expression was harvested by centrifuged at 5,000 rpm at 4°C for 15 min. The cell pellets were collected and then stored at -80°C for at least 2 hours.

The cell pellets were re-suspended in the cell lysis buffer (50 mM Na_2_PO_4_, 300 mM NaCl, 10 mM imidazole, 1 mM MgCl_2_, 2 mM dithiothreitol (DTT), 0.03% Triton-X, pH 7.7) with the addition of Complete EDTA-free protease inhibitor cocktail (Roche, Mannheim, Germany). The cell suspensions were sonicated for 4 cycles of 30 second on ice with a 1-minute rest interval. The cell lysate was then centrifuged at 5,000 rpm, at 4 °C for 15 min. The supernatant was transferred into 50 mL centrifuge tubes and incubated with DNase I (Roche, Mannheim, Germany) at a final concentration of 100 µg/mL on ice with low agitation for 1 hour. Cell lysates were then centrifuged at 5,000 rpm and 4°C for 15 minutes. The supernatant was collected as a crude extract.

The recombinant protein was purified by the His-tag at its C-terminus using affinity chromatography. Two of 3-mL His-spin columns (ThermoFisher) were equilibrated with 2 fold of resin bed volume of His-Binding buffer (50 mM Na_2_PO_4_, 300 mM NaCl, 10 mM imidazole, 1 mM MgCl, pH 7.7, Zymo) for 5 times. Six mL of crude extracts was loaded into each of the His-spin columns and incubated for 30 min at 4^°^C with low agitation. The columns were then centrifuged at 700 *X* g for 2 min at 10 °C. The loading steps were repeated until all crude extracts were loaded into the columns. The columns were then washed three times with Wash buffer (50 mM Na_2_PO_4_, 300 mM NaCl, 50 mM imidazole, 1 mM MgCl, pH 7.7, Zymo) to remove unwanted protein. The recombinant protein was then eluted with Elution buffer (50 mM Na_2_PO_4_, 300 mM NaCl, 250 mM imidazole, 1 mM MgCl, pH 7.7, Zymo). The elution fractions <u>containing enzyme activity</u> were pooled and dialyzed against 50 mM HEPES buffer, pH 7.4. Protein concentrations were quantified by the Bradford assay, using BSA as a standard. Western blot analysis was performed to determine the purity of the target protein. The pools containing recombinant nPA protein were analyzed by 4-20% SDS-PAGE and then transferred to PVDF membrane. The membrane was blotted with *Influenza* A PA Polyclonal antibody (1:20,000) as the primary antibody. The HRP-conjugated antirabbit IgG was used to attached against the primary antibody. The blot was developed with HRP substrates and the image was taken by iBright CL750 Imaging System instrument (Invitrogen).

### 2.5 *In vitro* endonuclease activity assay

A fluorescence resonance energy transfer (FRET)-based endonuclease assay (Figure 1) was employed and validated to assess the functionality of the recombinant nPA protein as described in previously published method ^13^ with modification. A FRET-based endonuclease assay was set up in Black Costar 96-well microplates. Dual-labeled single-strand DNA oligonucleotide with FAM (fluorophore) and a nonfluorescent quencher conjugated to its 5’ and 3’ terminals, respectively (5’ – 6FAM GGTTCCATGCTATATCTGGGACC MGBNFQ – 3’), synthesized by Thermo Fisher Scientific, was used as substrate (the 23mer ssDNA substrate described in the Figure 1). In a 100 µL reaction, nPA protein of the concentrations (25, 12.5, 6.25, 3.125, and 1.56 µg/mL) of a serial 1:2 dilution was added to the reaction buffer containing 50 mM HEPES buffer (pH 7.8), 150 mM NaCl, and 1 mM MnCl_2_, along with 200 nM substrate. Wells containing all assay components except the protein were used as background control. The fluorescence signals would be released due to the endonuclease cleavage and measured in BioTek Synergy Neo2 microplate reader at 39 s intervals, over 60 min at 37°C (*λ*_ex_ = 485 nm, *λ*_em_ = 535 nm). The gain was set to 50, and the fluorescence emission read for each concentration was background corrected. The slope of relative fluorescence unit (RFU) was plotted against the protein concentration to determine the optimum concentration of nPA protein applied in the endonuclease inhibitory assay.

**Figure 1.**
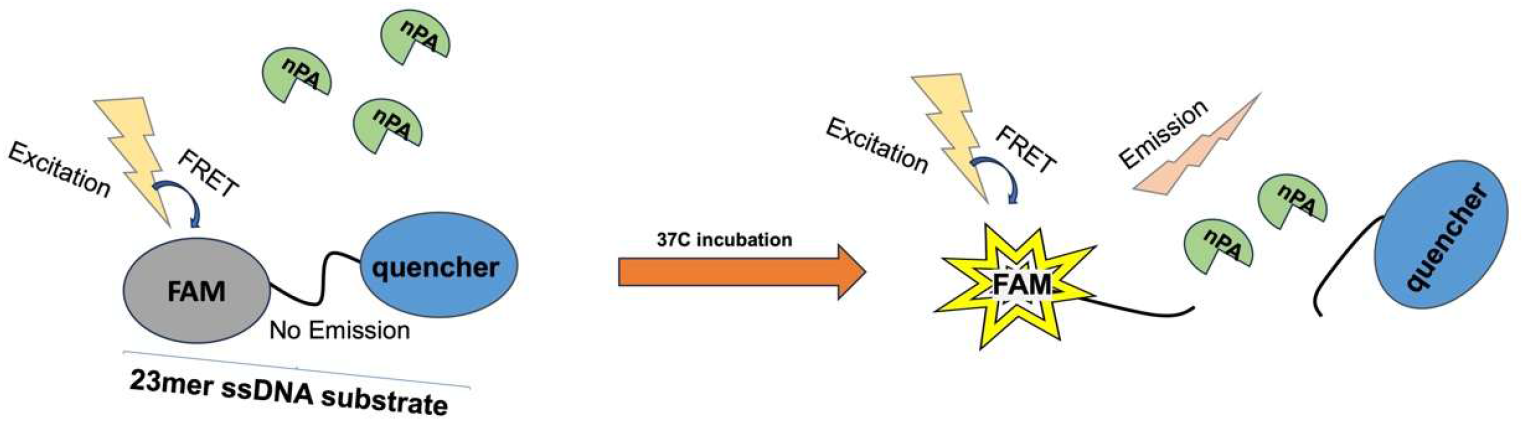
FRET-based endonuclease assay scheme.

The FRET-based endonuclease assay was performed to evaluate the antiviral property of the juices of 21 American elderberry cultivars and 1 European elderberry (EEB) cultivar. For sample preparation, 1.5 mL of juice from each elderberry cultivar was centrifuged for 5 min at 10,000 rpm and 10 °C to remove any leftover pomace and precipitate from the juice. The supernatant was transferred to a clean 1.5 mL Eppendorf tube for testing. Ten microliters of the samples were added to the well followed by the reaction buffer (50 mM HEPES buffer pH 7.8, 150 mM NaCl, and 1 mM MnCl_2_) and 200 nM substrate in 96-well black microplates. Then, 25 ng/µl of nPA protein was added to each well. Wells containing all assay components except the protein were used as background control. The plate was then placed in the plate reader and run the course of the assay at 37°C for 60 minutes. Selected bioactive compounds of American elderberry were screened for antiviral activity. Bioactive compounds were dissolved in pure DMSO and serially diluted to 20, 10, and 5 µM. An endonuclease inhibitory assay was carried out using the same method described above. Gallic acid, a known inhibitor of influenza endonuclease activity from a natural product ^14^, was used as a positive control in this assay. Wells containing all assay components except the protein were used as background control. Wells containing all assay components except the compounds were used as a negative control. Percent of inhibition of each compound was determined by normalizing the slope of fluorescence emission reads to that of the positive and negative controls.

### 2.6 Statistical analysis

The results of the analysis were expressed as means ± SEM (n=2). One-way analysis of variance (ANOVA) with Tukey’s multiple comparison test was performed using the R programming language version 4.5.1 to determine significant differences among mean values. Statistical significance was defined at *p-*value of < 0.001.

## 3. Results

The *npa gene* (Figure 2) has been successfully cloned into the T7 expression vector pET303 as confirmed by sequence analysis at the MU DNA core facility. The *npa gene* cloned pET303 was subsequently transferred into the expression host *E. coli* BL21(DE3) RIL strain (Figure 3A). The recombinant *n*PA protein was expressed in a 2-L culture and then isolated by the His-tag using 6xHis-tag affinity chromatography. Western blot analysis of the purified protein revealed a major band of protein species at ∼30 kDa that is the estimate molecular weight of *n*PA protein, confirming the successful expression and purification (Figure 3B, red arrow).

**Figure 2.**
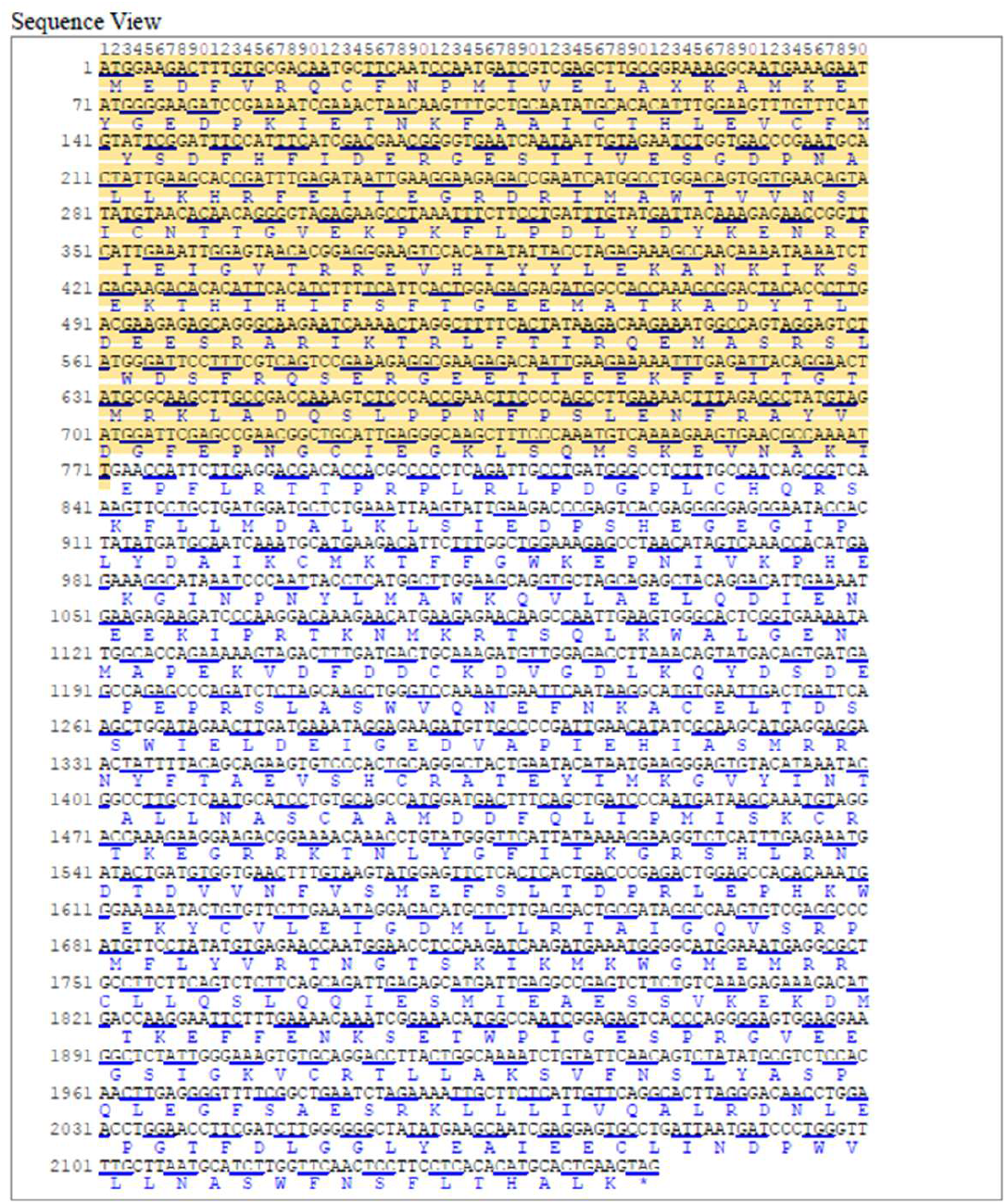
*Influenza* A (A/California/07/2009(H1N1)) segment 3 polymerase PA (PA) gene, complete cds (Source: NCBI; NC_026437.1). N-terminal domain of *PA gene* (1-771 bp) is highlighted in yellow.

**Figure 3.**
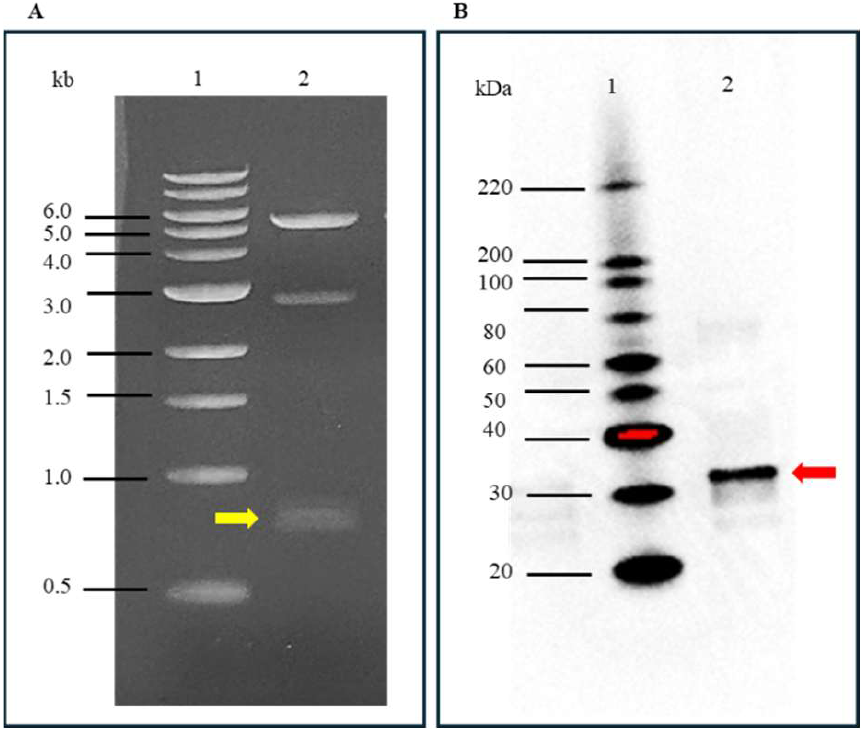
A. Analysis of *nPA* clones (yellow arrow) after XbaI and XhoI restriction enzyme digestion on 1% agarose gel electrophoresis. Lane 1, 1kb DNA ladders; lane 2, *npa* cloned pET303 isolated from BL21(DE3) RIL. B. Isolated recombinant *n*PA (red arrow) separated by 4-20% SDS-PAGE, transferred to PVDF membrane and developed against *Influenza* A PA Polyclonal antibody (1:20,000) as the primary antibody. The HRP-conjugated anti-rabbit IgG was used to attach to the primary antibody. The blot was developed with HRP substrates, and the image was taken by iBright CL750 Imaging System instrument (Invitrogen). Lane 1. protein ladder; lane 2. recombinant *n*PA protein (Size = 30.58 kDa).

An endonuclease assay method to determine the enzymatic activity of the recombinant nPA protein was established and validated using a FRET-based approach. The fluorescence intensity was recorded in a time course to monitor the cleavage activity of the nPA protein using the single-stranded DNA (ssDNA) substrate. The endonuclease activity of Recombinant nPA protein at varying concentrations was represented by fluorescence intensity RFU in Figure 4A. Enzyme activity was determined from the slope of fluorescence intensity over time using linear regression analysis. The resulting slopes were plotted against nPA protein concentration, demonstrating a positive correlation between enzyme concentration and endonuclease activity (Figure 4B). Protein concentration of 25 µg/ml was chosen for the inhibitory assay. To validate this endonuclease assay method for evaluating the inhibitory potential of natural product-derived compounds, gallic acid was used as a model inhibitor. Gallic acid was tested at varying concentrations of 20, 10, 5, and 0 µM, and the results demonstrated a dose-dependent inhibition of nPA endonuclease activity (Figure 4C), reflected by a decrease in the slope of enzymatic activity (Figure 4D).

**Figure 4.**
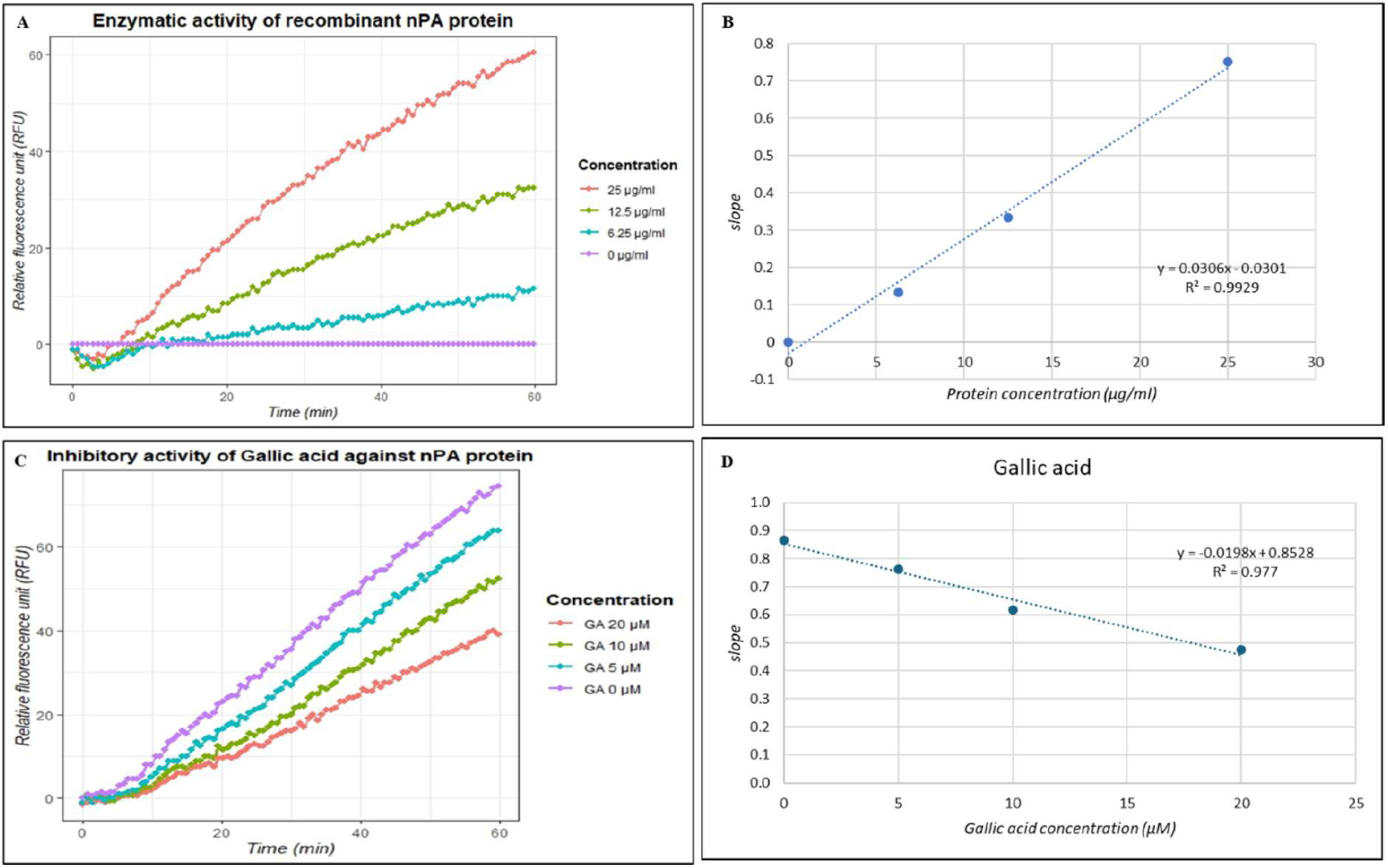
Detection of nPA endonuclease activity determined by FRET-based assay. The fluorescence intensity of each reaction was recorded at the indicated time points. (A) Endonuclease activity as a function of nPA protein concentration. The various concentration (25, 12.5, 6.25, and 0 µg/ml) of nPA protein was incubated with 200 nM ssDNA substrate and the fluorescence intensity was measured. (B) A concentration-dependent increase in fluorescence intensity over time, represented as the slope of the linear regression of nPA enzymatic activity. (C). Endonuclease activity as a function of Gallic acid inhibitor concentration. (D) Inhibitory effect of gallic acid at varying concentrations (20, 10, 5, and 0 µM) against nPA endonuclease activity (n=2) presented as the slope of nPA enzymatic activity.

**Figure 5.**
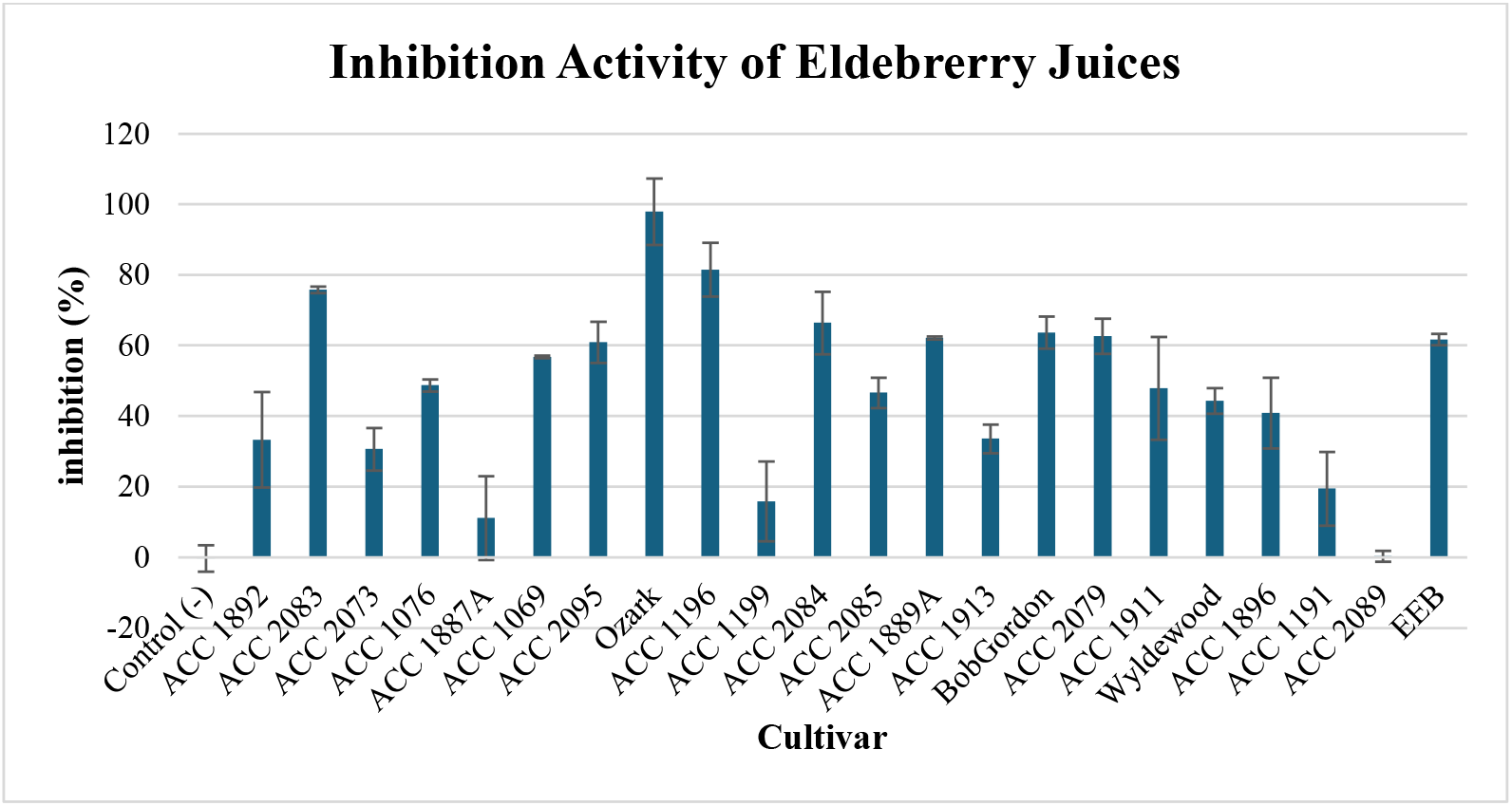
Inhibitory effects of elderberry juices from various cultivars on influenza nPA protein. The percentage of inhibitory effect as a function of the species of elderberry cultivars, determined by endonuclease inhibitory assay in duplicate. Data presented as mean ± SEM (n=2). [European elderberry cultivar Haschberg = EEB]

The antiviral activity of elderberry juices from 3 established cultivars and 18 wild accessions of American elderberry cultivars was evaluated using the validated endonuclease inhibitory assay against the recombinant nPA protein produced in this project. All of cultivar juices were tested as undiluted juices due to the complexity of the juice matrix. Results indicated that juices of American elderberry cultivars exhibited varying level of inhibition of nPA endonuclease activity, with juices from cultivar Ozark showing the strongest inhibitory effect, followed by accession 1196 and 2083 with percent inhibition of 97.86% ± 9.42, 81.48% ± 7.68, and 75.73% ± 0.93, respectively. Among the American elderberry cultivars, Bob Gordon, and wild accessions 2084, 2079, 1889A, and 2095, demonstrated moderate inhibition levels of approximately 60%, with percent inhibition of 63.61% ± 4.55, 66.41% ± 8.86, 62.67% ± 4.98, 62.09% ± 0.45, 61.66% ± 1.59, and 60.88% ± 5.86, respectively. Wild accession 1191, 1199, and 1887A showed low inhibition levels below 25%, with inhibition levels of 19.42% ± 10.47, 15.83% ± 11.30, and 11.11% ±11.89, respectively. Interestingly, wild accession 2089 did not show the inhibitory effect.

In order to compare the antiviral activity of American elderberry with European elderberry, the juice of the European elderberry cultivar Haschberg (EEB) was also tested and exhibited the level of 61.65% ± 1.59 inhibitory effect. The results showed that four of American elderberry juices outperformed European elderberry juice in inhibitory potential to nPA endonuclease in this assay.

To further evaluate the possible bioactive compounds responsible for the antiviral activity of American elderberry juices, 32 bioactive compounds previously identified in American elderberry juices were tested using the endonuclease inhibitory assay. These compounds, which included anthocyanins, flavonoids, and other polyphenolic compounds, were selected based on their relevance to American elderberry’s unique phytochemical profile and were previously evaluated for their bioactive potential ^15^. Testing these compounds provided a more direct assessment of elderberry’s ability to inhibit the *n*PA endonuclease activity of *influenza* A virus, hence providing the connection between metabolomic findings and functional antiviral validation.

Among the 32 compounds tested, eight compounds showed a dose-dependent inhibition against nPA protein. Each compound was evaluated at three different concentrations (20, 10, and 5 µM), and the half maximal inhibitory concentration (IC_50_) values were calculated based on the linear regression of the slope of their inhibitory activity. This analysis enabled a quantitative comparison of the inhibitory potency of each compound tested (Figure 6). Two compounds exhibiting the strongest inhibitory activity were classified as phenolic acid (gallic acid) and flavonoids (myricetin). Gallic acid showed the strongest inhibition, with an IC_50_ value of 11.56 ± 1.80 µM, followed by myricetin with an IC_50_ value of 16.02 ± 0.06 µM. Caffeic acid, a phenolic acid, and luteolin, a flavonoid, showed moderate inhibition with IC_50_ values of 22.75 ± 0.9 µM and 27.92 ± 0.72 µM. Other phenolic acids, such as chlorogenic acid and neochlorogenic acid, also displayed inhibitory effects, although with lower potency (IC_50_ values of 33.28 ± 2.664 and 49.68 ± 4.45, respectively). In contrast, most anthocyanins, such as cyanidin-3-*O*-sambubioside and cyanidin-3-*O*-rutinoside, showed IC_50_ values higher than 50 mM, suggesting that flavonoids and phenolic acids may be the primary contributors to the antiviral potential of elderberry (Table 1).

**Figure 6.**
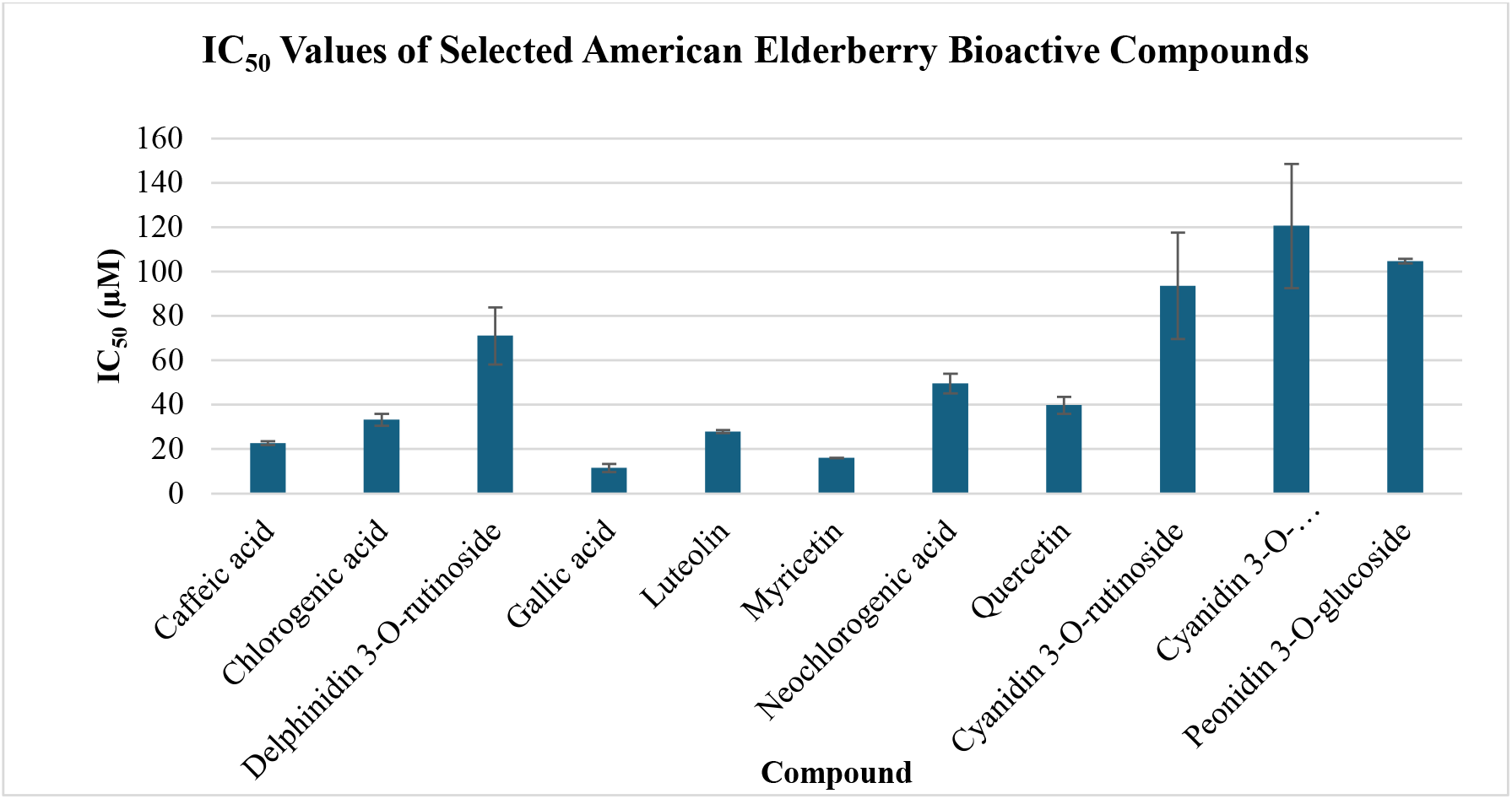
The half maximal inhibitory concentration (IC_50_) values of American elderberry compounds against *Influenza* A *n*PA protein. Data presented as mean value ± SEM (n=2)

## 4. Discussion

In this study, we utilized a FRET-based endonuclease assay to evaluate the inhibitory activity of elderberry juices against the *Influenza* A PA protein. This assay has previously been used to screen novel bioactive compounds and shown to be reliable for determining the inhibitory activity of compounds ^13,12^. Our results demonstrated that American elderberry juices inhibited *n*PA endonuclease activity, with apparent differences across cultivars that likely reflect the variation in phytochemical composition. Notably, American elderberry juices of several cultivars Ozark, accession 1892, 1196, 2084 outperformed European elderberry (*S. nigra*, cultivar Haschberg). These findings suggest that American elderberry could have a more diverse and potentially superior chemical profile compared to its European counterpart, making it a better source of raw material for the production and development of antiviral supplement products.

Previous studies utilizing European elderberry extract as a source of antiviral agents have identified the major anthocyanin, cyanidin 3-glucoside, as a key compound responsible for antiviral activity by inhibiting the viral hemagglutinin glycoprotein, thereby preventing viral attachment to host cells ^5^. However, our findings suggested that different classes of metabolites, including flavonoids and phenolic acids, from American elderberry contribute to the antiviral activity against *Influenza* A virus through different mechanisms of action.

In addition to the inhibitory effects observed in the juices, several compounds were found to inhibit *Influenza* PA endonuclease activity, including flavonol compounds, gallic acid, myricetin, and luteolin. The inhibitory activity of gallic acid against influenza PA endonuclease has previously been reported in studies investigating the bioactive compounds of green tea ^14^. Interestingly, three isomers, chlorogenic acid, cryptochlorogenic acid, and neochlorogenic acid, demonstrated various levels of inhibition against *n*PA endonuclease activity. This finding suggested potential structural specificity in the interaction between the inhibitor and the active site of the PA N-terminal domain. However, further structural binding analysis is needed to elucidate the precise molecular mechanism underlying this selectivity. In contrast, several anthocyanin compounds that are unique to American elderberries didn’t inhibit the *n*PA endonuclease activity (Table 2).

Aside from that individual elderberry compounds exhibit inhibitory effects, potential synergistic interactions among multiple metabolites may lead to enhanced antiviral potency. Further studies are needed to elucidate how a combination of multiple metabolites could be optimal to the overall antiviral activity of elderberry. Nevertheless, these findings not only support the inhibitory effects observed at the juice level but also identify chemical scaffolds that can be used in the nutraceutical industry. This study provides the scientific evidence that American elderberry juices and their metabolite constituents interfere with the function of the *Influenza* nPA protein.

## 5. Conclusion

Elderberry exhibits potent antiviral properties against *Influenza*, acting through inhibition of viral replication mechanisms. This study demonstrated, for the first time, the inhibitory effect of elderberry juice against the nPA protein. The juice of several American elderberry cultivars outperformed the inhibitory potential of European elderberry juice, with the American elderberry cultivar Ozark showing the most potent inhibitory effect. Elderberry compounds gallic acid, myricetin, caffeic acid, and luteolin showed the most potent inhibitory effect against *n*PA endonuclease activity *in vitro*. In contrast, anthocyanin compounds didn’t inhibit the *n*PA endonuclease activity. This finding supports the use of American elderberry as a superior raw material for application in nutraceutical formulations that aim at developing a natural product supplement to help manage Influenza symptoms.

## Acknowledgments

This work is supported by the USDA-NIFA-SCRI 2021-51181-35860, “Moving American Elderberry into Mainstream Production and Processing”. We would like to acknowledge the USDA Specialty Crop Research Initiative (SCRI), the Missouri Department of Agriculture (MDA) Specialty Crop Block Grant Program (SCBGP), the Center for Agroforestry at the University of Missouri, and USDA/ARS Dale Bumpers Small Farm Research Center under agreement number 58-6020-6-001 from the USDA Agricultural Research Service for their support.

